# Epidemiology of drug-resistant tuberculosis in Chongqing, China: a retrospective observational study from 2010 to 2017

**DOI:** 10.1101/610048

**Authors:** Bo Wu, Ya Yu, Changting Du, Ying Liu, Daiyu Hu

## Abstract

China is one of the top 30 countries with high multidrug-resistant tuberculosis (MDR-TB) and rifampin-resistant tuberculosis (RR-TB) burden. Chongqing is a southwest city of China with a large rural population. A retrospective observational study has been performed based on routine tuberculosis (TB) surveillance data in Chongqing from 2010 to 2017. The MDR/RR-TB notification rate increased from 0.03 cases per 100,000 population in 2010 to 2.09 cases per 100,000 population in 2017. The extensively drug-resistant TB (XDR-TB) notification rate has increased to 0.09 cases per 100,000 population in 2017. There was a decreasing detection gap between the number of notified MDR/RR-TB cases and the estimate number of MDR/RR-TB cases among all notified TB cases. The treatment success rate of MDR/RR-TB was 50.66% in this period. The rate of MDR/RR-TB in new TB cases was 6.23%, and this rate in previously treated TB cases was 32.7%. Despite the progress achieved, the prevalence of MDR/RR-TB was still high facing challenges including detection gaps, the regional disparity, and the high risk for MDR/RR-TB in elderly people and farmers. Sustained government financing and policy support should be guaranteed in the future.

## Introduction

Drug-resistant tuberculosis (DR-TB) threatens global TB prevention and control. MDR-TB, defined as TB resistant to at least rifampin (RFP) and isoniazid (INH), has longer treatment regimens and lower treatment success rate with supplies of less effective and more toxic second-line drugs [1]. RR-TB, defined as TB resistant to at least RFP, also requires a second-line regimen, and 83% of RR-TB cases have MDR-TB [2].

From 2006 to 2014, China had implemented a project targeting MDR-TB through a partnership with the Global Fund to Fight AIDS, Tuberculosis, and Malaria (Global Fund) [3]. In 2009, the Bill and Melinda Gates Foundation and the Chinese Ministry of Health (now called the National Health Commission of China) announced the initiation of a comprehensive MDR-TB development programme aiming to provide innovative MDR-TB diagnosis tool, improve affordability of the treatment and the treatment success rate. A health system reform has also been launched since 2009 in China [4]. However, the prevalence of MDR-TB remains high. China is one of the 30 high MDR/RR-TB burden countries, and has an estimated 73,000 incident cases of MDR/RR-TB in 2016, accounting for 12.15% of all MDR/RR-TB burden in the world [2].

Chongqing is a municipality directly under the central government in the southwest of China with about 30 million people. Approximately 23,000 TB patients were reported in Chongqing in 2017, and the TB notification rate was 77.1 cases per 100,000 population which was 27.5% higher than the rate of the whole country. In 2009 and 2013, Chongqing participated in the programme of the Bill and Melinda Gates Foundation and the Global Fund respectively. Significant improvement in MDR/RR-TB notification, diagnosis and treatment has been achieved since 2009. In 2017, 643 MDR/RR-TB cases were notified in Chongqing, and the MDR/RR-TB notification rate is 2.1 cases per 100,000 population. In 2017, 29 cases with XDR-TB, defined as MDR-TB plus additional resistant to a fluoroquinolone and a second-line injectable, were notified in Chongqing. However, gaps between MDR/RR-TB notifications and the estimate number of MDR/RR-TB cases remains big. In 2017, there were about 1000 MDR/RR-TB cases undetected in Chongqing.

This observational retrospective study tried to explore the trend of MDR/RR-TB and the changes of drug resistance patterns based on routine surveillance data in Chongqing from 2010 to 2017. Our study may serve as a reference for the future policy of MDR/RR-TB control in Chongqing and other regions with similar situations.

## Methods

### Study design and data collection

This was an observational retrospective study of notified MDR/RR-TB cases from 2010 to 2017 in Chongqing. The information of MDR/RR-TB cases came from the national electronic TB surveillance system.

The DR-TB notification rate from 2010 to 2017 was analyzed, including MDR/RR-TB and XDR TB. The MDR/RR-TB cases were stratified by age, sex, and occupation for trend analysis. The information of sex, age and occupation of MDR/RR-TB cases came from the national TB surveillance system. The socio-demographic data came from Chongqing Statistical Yearbook from 2010 to 2017. According to the Chongqing Statistical Yearbook, the population in Chongqing was divided into four age groups: 0-17 years, 18-34 years, 35-59 years, and over-60 years. The occupation of MDR/RR-TB cases was divided into farmers and non-farmers. The regional disparity of TB notification has also been evaluated. According to the division of local government, there were four regions in Chongqing including: the Urban Districts, New Urban Development Districts, Northeast Districts and Southeast Districts. These regions were different in terms of socioeconomic development.

The detection gap between the number of notified MDR/RR-TB cases and the estimated number of MDR/RR-TB cases among all notified TB cases has been evaluated. According to the most recent measured MDR/RR-TB proportion in new and previously treated TB cases from the national DR-TB survey in 2013, the number of MDR/RR-TB cases among all notified TB cases could be estimated. This proportion could be used to multiply the number of notified TB cases, and the estimated number of MDR/RR-TB cases could be calculated.

The information of MDR/RR-TB screening, DST, diagnosis, treatment, management and outcome has been recorded in the national electronic surveillance system since 2010. The data of MDR/RR-TB cases from 2010 to 2017 came from this system. Reporting of extra-pulmonary TB was not mandatory in this system, thus all data were based on the pulmonary MDR/RR-TB cases. Population and socioeconomic data came from Chongqing Statistical Yearbook from 2010 to 2017.

### Risk factor analysis

To explore risk factor associated with MDR/RR-TB in Chongqing, we have analyzed potential risk factors including age, sex, occupation, region, and treatment history. All MDR/RR-TB screening data from 2010 to 2017 were analyzed, and MDR/RR-TB cases have been compared with drug-susceptible TB cases.

### Drug resistance patterns

Drug susceptible test (DST) include both phenotypic (conventional) and genotypic (molecular) testing methods. In Chongqing, conventional DST was performed at the provincial reference laboratory using the proportion method on acid-buffer Lowenstein-Jensen (L-J) Medium. The first-line drugs tested for drug-resistance included INH, RFP, ethambutol (EMB), and streptomycin (SM). There were two second-line drugs reported, including ofloxacin (OFX) and kanamycin (KM). Rapid molecular test was performed at capable county-level laboratories and prefectural laboratories. In Chongqing, the main rapid molecular test included Xpert MTB/RIF and fluorescence PCR melting curve method. Both rapid molecular tests could detect INH and RFP resistance in one day. External quality assessment has been conducted through national reference laboratory of China.

According to definitions and reporting framework for tuberculosis of World Health Organization (WHO) [5], the treatment outcome from 2010 cohort to 2015 cohort was also analyzed. Treatment success was the sum of cured and treatment completed.

### Detection and treatment procedure

In Chongqing, the MDR/RR-TB control system was constructed of three levels: province, prefecture, and county. Provincial TB prevention and control institution was responsible for administration, supervision, DST, and surveillance.

There were 38 counties and districts in Chongqing, every one of which had a county-level designated clinic for TB. The county-level designated clinics detected TB cases according to the tuberculosis diagnostic criteria WS288-2008 of China [6], and screened MDR/RR-TB.

In early stages, inadequate resources limited the systematic development of screening. Screening had been conducted spontaneously in all counties and districts before 2010. With the introduction of some projects, screening was becoming more and more standardized. From March 2010 to December 2010, six counties and districts screened MDR/RR-TB supported by a domestic project, including Wanzhou district, Kaizhou district, Liangpin county, Yunyang county, Zhong county and Fengjie county. MDR/RR-TB suspects stemmed from sputum smear positive TB cases. From March 2011 to March 2012, five counties and districts screened MDR/RR-TB suspects from sputum smear positive TB cases supported by the Bill and Melinda Gates Foundation, including Jiangjin district, Dazhu district, Rongchang district, Tongliang district, and Yongchuan district.

Since 2013, screening in all counties and districts has supported by local government investment and the Global Fund. From 2013 to 2015, MDR/RR-TB suspects came from five high-risk groups, including (1) re-treatment cases with treatment failure or sputum smear positive cases with previous irregular treatment, (2) close contacts of MDR/RR-TB with sputum smear positive, (3) recurrence cases, (4) new cases persisting sputum smear positive at the end of 2^nd^ month after treatment, and (5) new cases of initial treatment failure. In 2016, 20% of new cases with sputum smear positive and five high-risk groups were screened. In 2017, all new cases and five high-risk groups were screened. A new TB case was defined as a patient who has never been treated or been treated less than 1 month, a re-treatment or previously treated case was defined as a patient who received TB treatment for 1 month or more previously, and a recurrence case was defined as a patient who was cured but diagnosed as TB again [5,7].

Since 2010, eighteen county-level laboratories had been furnished with rapid molecular test equipment, and seven prefectural laboratories had been set up. In the counties and districts furnished with rapid molecular test equipment, rapid molecular test was conducted firstly. If the result was resistant to INH and RFP or resistant at least to RFP and the suspect was part of five high-risk groups, the suspect would be diagnosed as MDR/RR-TB. If the first result was positive and the suspect was not part of five high-risk groups, another rapid molecular test would be required. Only when both results were positive, the suspect would be diagnosed as MDR/RR-TB. After rapid molecular test, the MDR/RR-TB cases would be referred to prefectural designated hospitals for treatment, and sputum culture was conducted in a county-level laboratory or a prefectural laboratory. If a positive culture result was confirmed, the strain of MDR/RR-TB suspect would be sent to the provincial reference laboratory for strain identification and DST. The results of strain identification and DST would be sent back to the county-level designated clinics and prefectural designated hospitals.

In the counties and districts without rapid molecular test equipment, sputum smear and culture was conducted in the county-level laboratories firstly. If the result of sputum culture was positive, the strain of MDR/RR-TB suspect would be sent to the prefectural laboratories for rapid molecular test. After rapid molecular test, the rest procedure was the same as the previous diagnosis and treatment process.

There were eight prefectural designated hospitals appointed by the provincial health administration, which were in charge of MDR/RR-TB diagnosis, hospitalization, and return visits in their local area. After diagnosis by a prefectural expert review committee, most MDR/RR-TB cases were enrolled into a standardized regimen. MDR/RR-TB cases stayed 2 months in the prefectural designated hospital, and then received an ambulatory treatment course for 22 months managed by the county-level center for disease control and prevention (CDC) or county-level TB prevention and control institution.

### Ethics approval

The study was approved by the Ethics Committee of the Institute of Tuberculosis Prevention and Control, Chongqing, China. In this study, there was no access to individual information. A secondary analysis based on reported data has been conducted and informed consent from individuals was not required. All methods were performed in accordance with relevant guidelines and regulations.

### Statistical analysis

The chi-square test for linear trend was used to test the trend of the notification rate. A logistic regression model was performed to assess association between notification and risk factors. A two-sided P-value less than 0.05 was taken as statistically significant. Statistical analyses were performed with the SPSS 22.0 software (SPSS, Inc., Chicago, IL, USA).

## Results

### Trend of MDR/RR-TB notification

From 2010 to 2017, 1,908 MDR/RR-TB cases were notified in Chongqing. Among them, 4.82% were XDR-TB cases. The MDR/RR-TB notification rate increased significantly from 0.03 cases per 100,000 population in 2010 to 2.09 cases per 100,000 population in 2017 (χ^2^ trend=1,623.15, *P*<0.05) (Fig 1). The XDR-TB notification rate increased significantly from 0.02 cases per 100,000 population in 2003 to 0.09 cases per 100,000 population in 2017 (χ^2^ trend=27.06, *P=*1.98×10^−7^).

**Fig 1.**
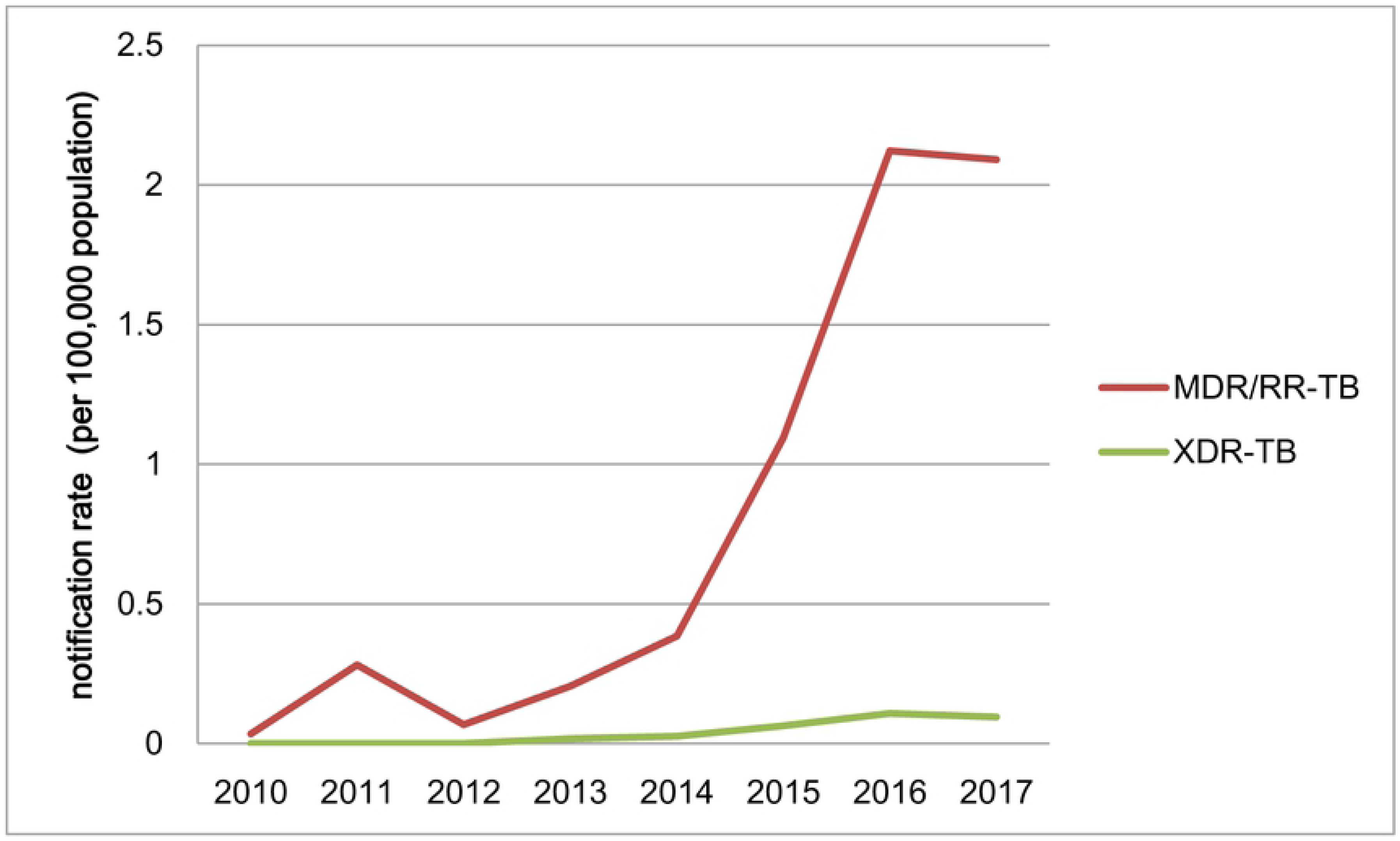
The trend of DR-TB notification rate from 2010 to 2017.

### MDR/RR-TB notification stratified by sex, age, and occupation

From 2010 to 2017, 70.02% of MDR/RR-TB cases were male. The MDR/RR-TB notification rate of male was significantly higher than the rate of female (χ^2^=2601.49, *P*<0.05). MDR/RR-TB notification rate of male increased significantly from 0.04 cases per 100,000 population in 2010 to 2.61 cases per 100,000 population in 2017 (χ^2^ trend=1,133.68, *P*=1.58×10^−248^). MDR/RR-TB notification rate of female also increased significantly from 0.02 cases per 100,000 population in 2010 to 1.14 cases per 100,000 population in 2017 (χ^2^ trend=536.82, *P=*9.26×10^−119^).

From 2010 to 2017, the trend of age-specific MDR/RR-TB notification rate increased significantly among all age groups. The age-specific MDR/RR-TB notification rate was significantly different in four age groups (χ^2^=2106.04, *P*<0.05). The MDR/RR-TB cases aged 18-34 and 35-59 accounted for 81.24% of all MDR/RR-TB cases. The MDR/RR-TB notification rate in the age group 18-34 years had a significant increase from 0.03 to 2.73 cases per 100,000 population (χ^2^ trend=585.3, *P=*2.63× 10^−129^). The MDR/RR-TB notification rate in the age group 35-59 years also increased from 0.06 to 2.27 cases per 100,000 population (χ^2^ trend=764.9, *P=*2.31× 10^−168^). The MDR/RR-TB cases aged 0-17 had the lowest notification rate, and the notification rate increased significantly from 0 to 0.3 cases per 100,000 population (χ^2^ trend=61.9, *P*=3.62× 10^−15^). The proportion of the MDR/RR-TB cases aged 0-17 was also lowest, which was 3.14%. The MDR/RR-TB notification rate of over-60-year age group increased significantly from 0 to 1.71 cases per 100,000 population (χ^2^ trend=264.69, *P*=1.63×10^−59^) (Fig 2).

**Fig 2.**
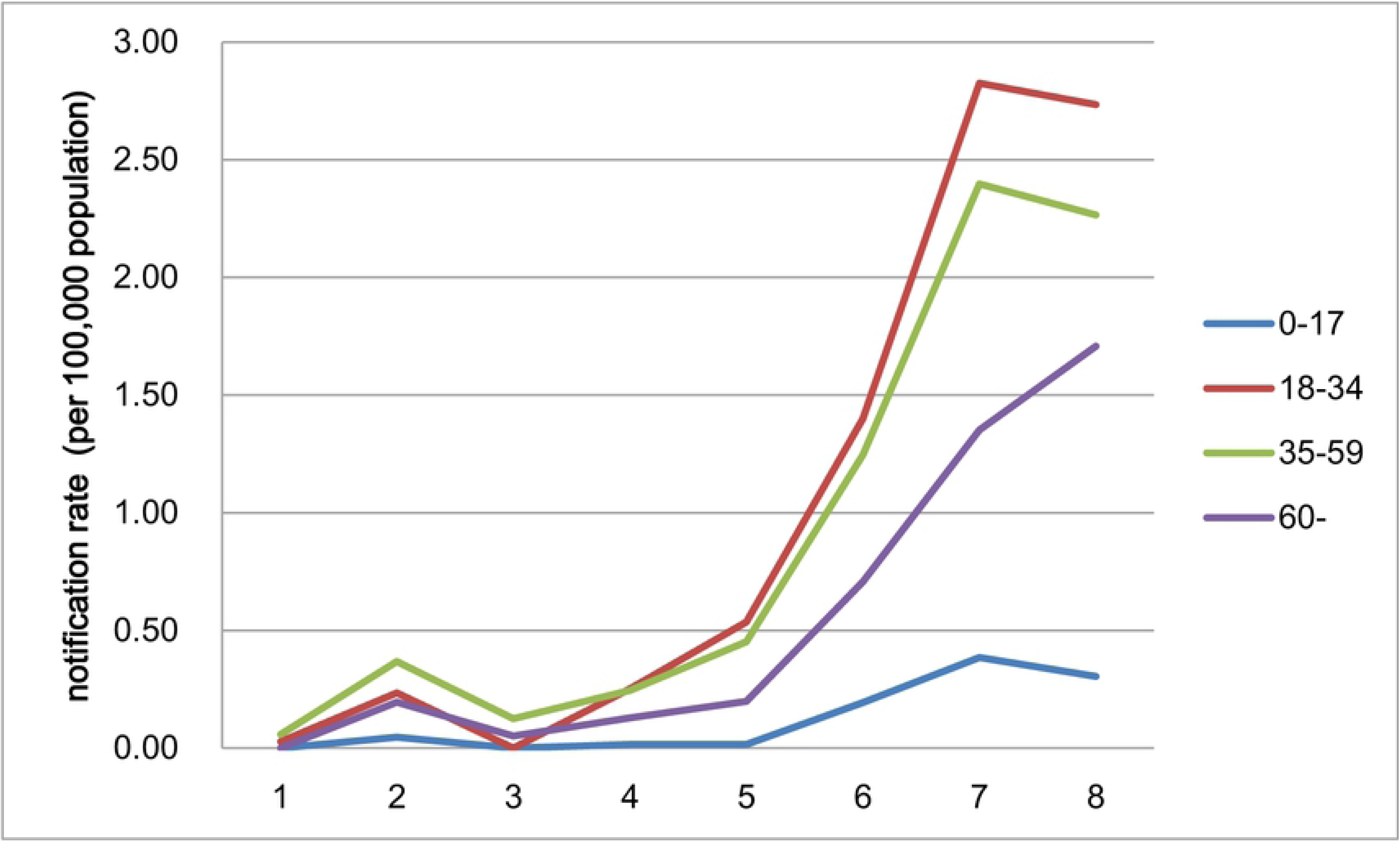
The MDR/RR-TB notification rate in different age groups from 2010 to 2017.

The MDR/RR-TB notification rate of farmers was significantly higher than the rate of all other occupations from 2010 to 2017 (χ^2^=2616.58, *P*<0.05). The MDR/RR-TB notification rate of farmers has significantly increased (χ^2^ trend=499.75, *P*=1.08×10^−110^).

### Regional disparity of MDR/RR-TB notification

The MDR/RR-TB notification rate was significantly different in four regions (χ^2^=2102.04, *P*<0.05), and the MDR/RR-TB notification rate of Urban Districts has been higher than other three regions since 2014(Fig 3). The MDR/RR-TB notification rate of Urban Districts has significantly increased from 0.16 to 7.72 cases per 100,000 population (χ^2^ trend=1,196.76, *P*=3.09× 10^−262^). In other three regions, The MDR/RR-TB notification rate was less than 1 case per 100,000 population.

**Fig 3.**
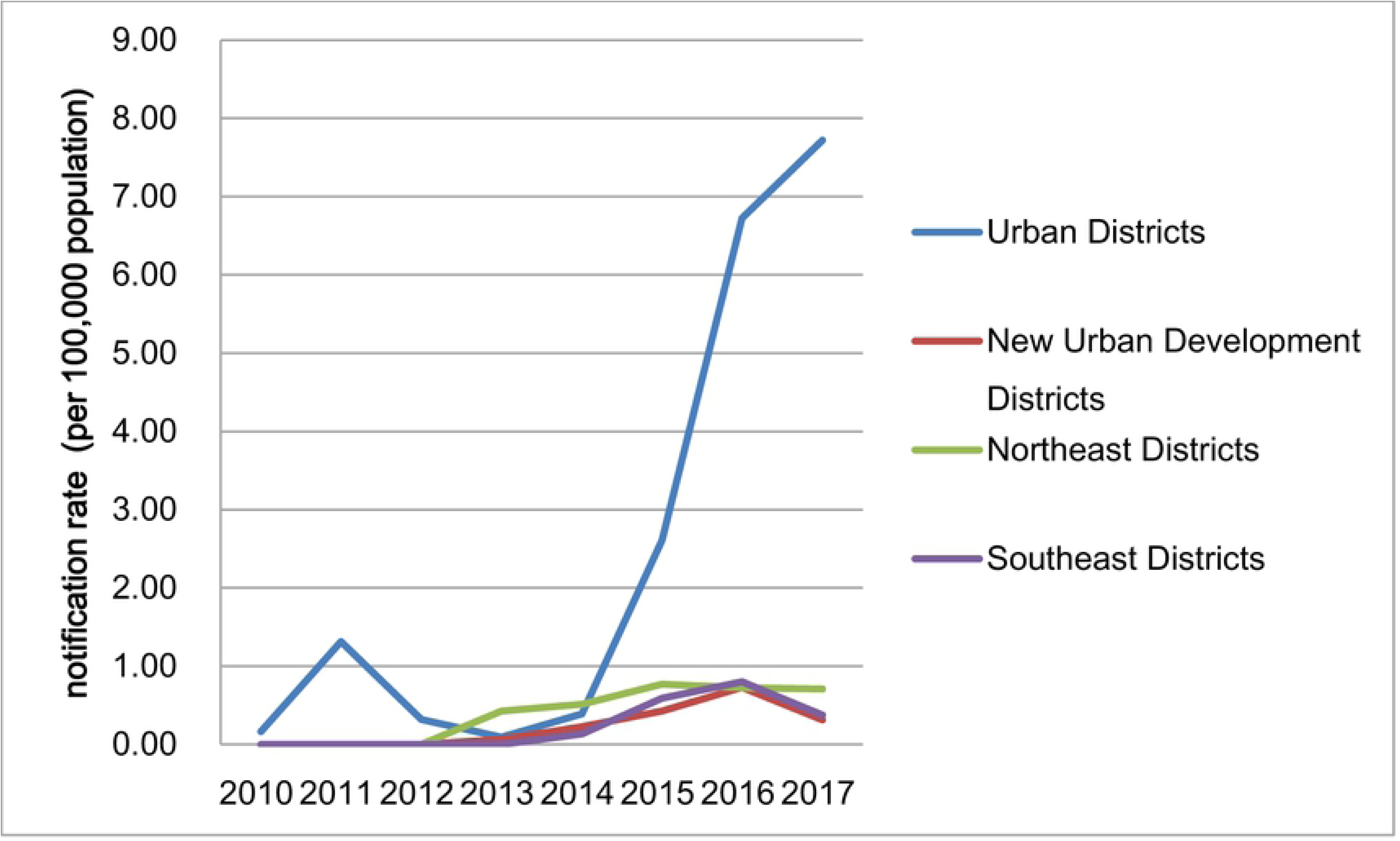
The MDR/RR-TB notification rate of different regions from 2010 to 2017.

### Detection gap

According to the national DR-TB survey in 2013, the MDR/RR-TB proportion in new TB cases was 7.1% and the MDR/RR-TB proportion in previously treated TB cases was 24%^2^. The estimated number of MDR/RR-TB cases among all notified TB case from 2010 to 2017 was 15104, which was 691.61% higher than the number of MDR/RR-TB cases found now. The detection gap between the number of notified MDR/RR-TB cases and the estimate number of MDR/RR-TB cases among all notified TB cases has significantly declined from 7.26 to 3.35 cases per 100,000 population (χ^2^ trend=1,446.45 *P*<0.05) (Fig 4).

**Fig 4.**
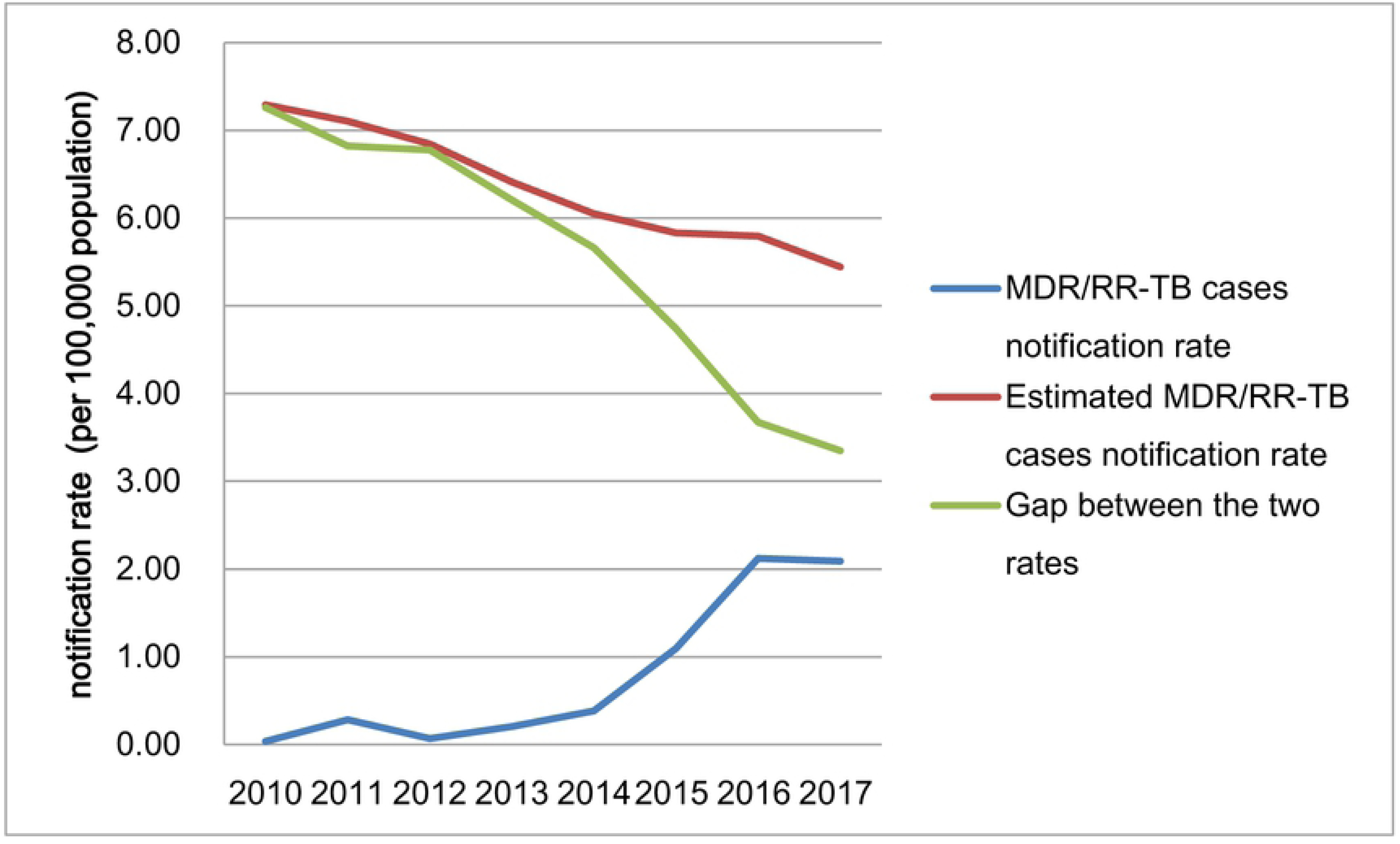
The notification gaps from 2010 to 2017.

### Treatment outcome

There were 618 MDR/RR-TB cases notified from 2010 to 2015, and the treatment success rate of MDR/RR-TB was 50.66% in this period. The death rate, treatment failure rate and rate of loss to follow-up were 13.49%, 9.21%, and 13.82% respectively. There were 132 cases who were not involved in treatment. In the 486 cases involved in treatment, 124 cases had not treatment outcome information reported. The treatment success rate of MDR/RR-TB increased significantly from 20% in 2011 cohort to 51.81% in 2015 cohort (χ^2^ trend=4.37, *P*=0.04) (Fig 5).

**Fig 5.**
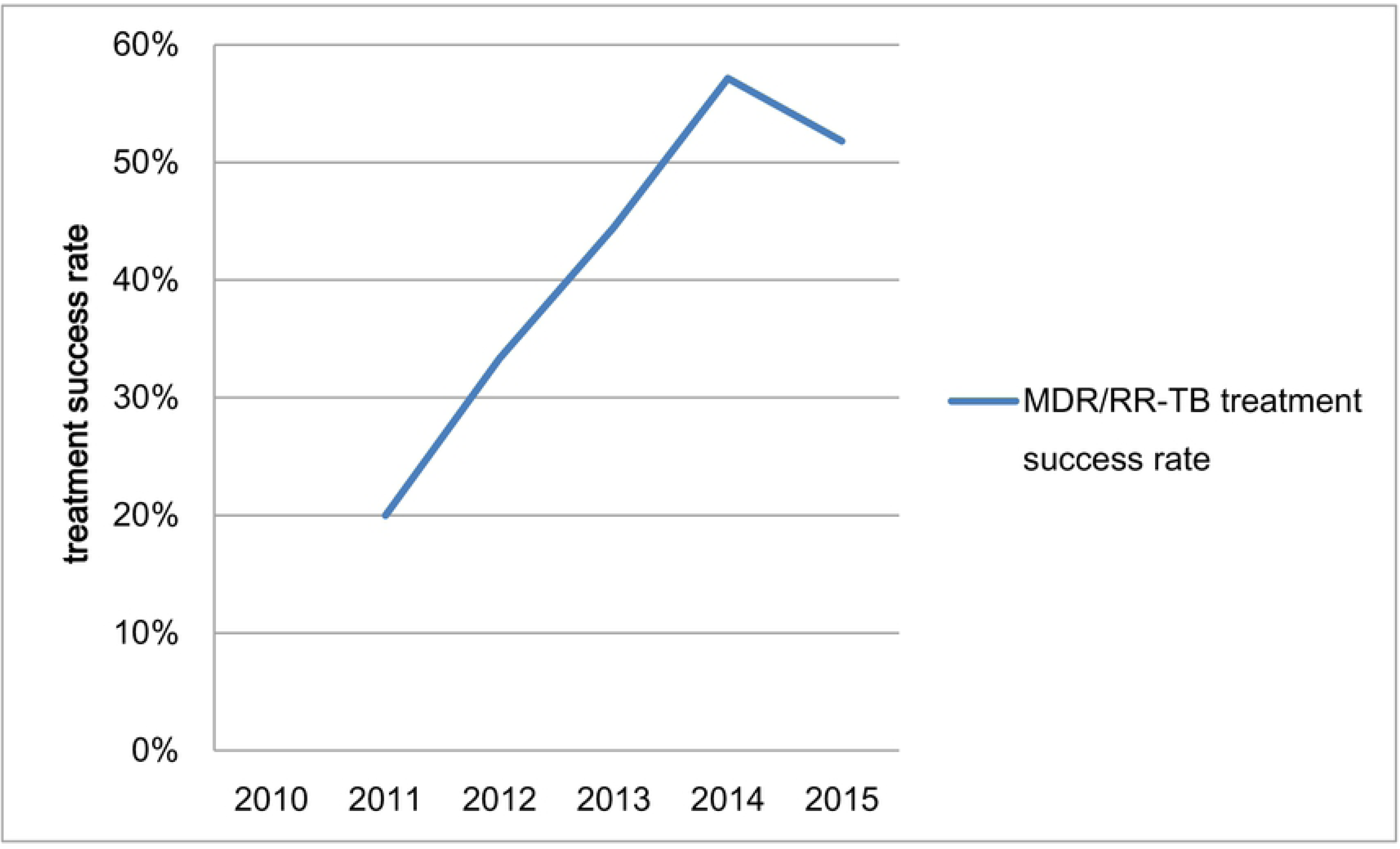
The MDR/RR-TB treatment outcome.

### Drug resistance patterns

From 2010 to 2017, the rate of drug resistance for RFP in new TB cases was 6.23%, and this rate in previously treated TB cases was 32.7%. Among all 12230 TB cases screened, RFP had the highest rate of drug resistance which was 17.13%, followed by INH (15.37%), SM (14.18%), EMB (9.43%), OFX (9.15%), and KM (2.42%). The drug resistance rate for RFP increased significantly from 3.07% in 2010 to 15.53% in 2017 (χ^2^ trend=172.92, *P*=1.71×10^−39^). The drug resistance rate for INH had a similar trend which increased significantly from 5.46% in 2010 to 12.48% in 2017 (χ^2^ trend=60.19, *P*=8.6× 10^−15^). The other four drugs also showed a significant rising trend, including EMB (χ^2^ trend=112.52, *P*=2.75× 10^−26^), SM (χ^2^ trend=65.55, *P*=5.66× 10^−16^), OFX (χ^2^ trend=31.37, *P*=2.14× 10^−8^), and KM (χ^2^ trend=11.47, *P*=0.001).

From 2016 to 2017, the drug resistance rate for RFP has declined significantly from 16.42% to 5.69% in new TB cases (χ^2^ trend=77.24, *P*=1.52× 10^−18^) (Fig 6). In 2016 and 2017, the drug resistance rate for RFP was 38.13% and 37.62% respectively in previously treated TB cases, which didn’t change significantly (χ^2^ trend=0.08, *P*=0.78) (Fig 7).

**Fig 6.**
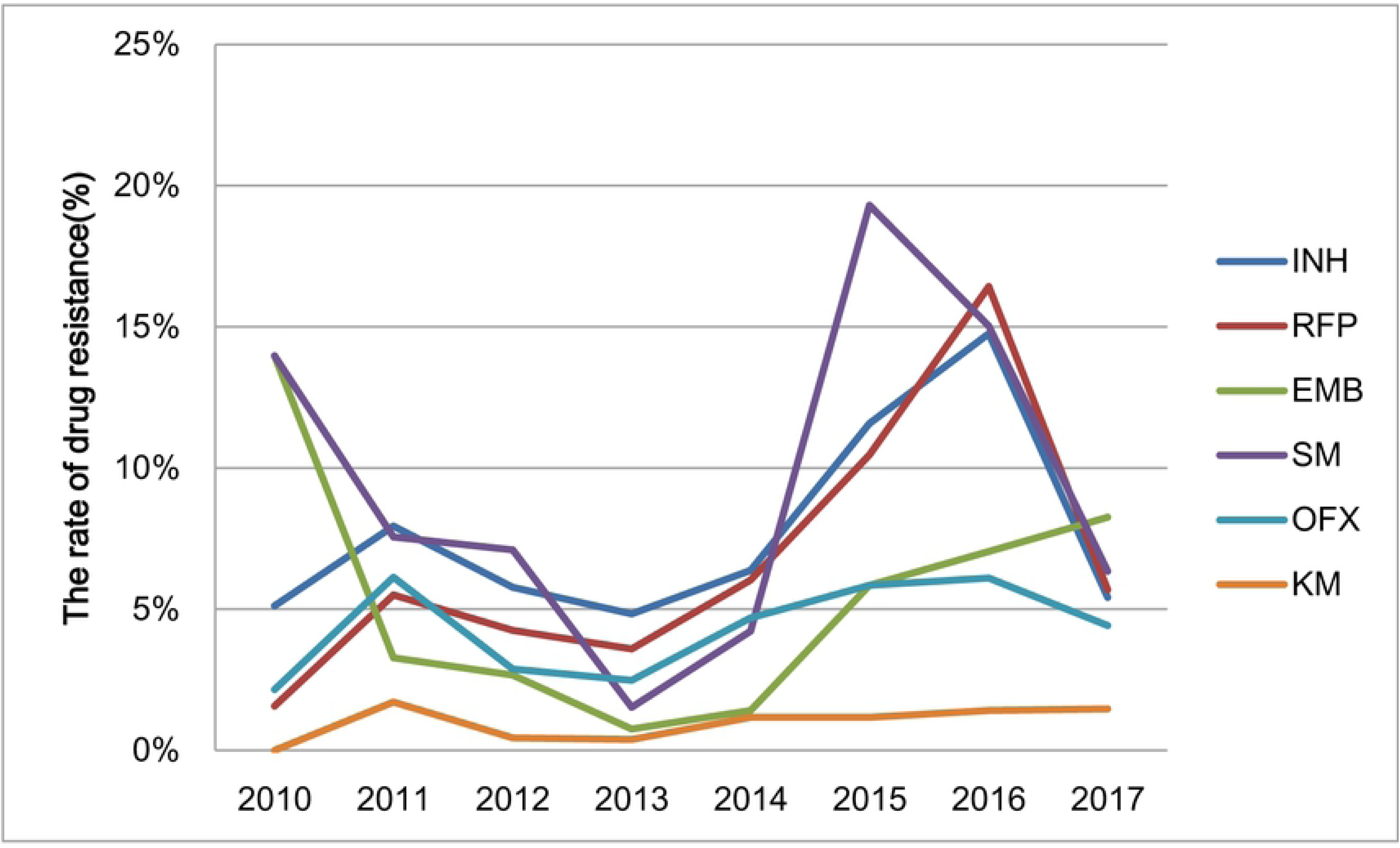
The drug resistance patterns in new TB cases from 2010 to 2017.

**Fig 7.**
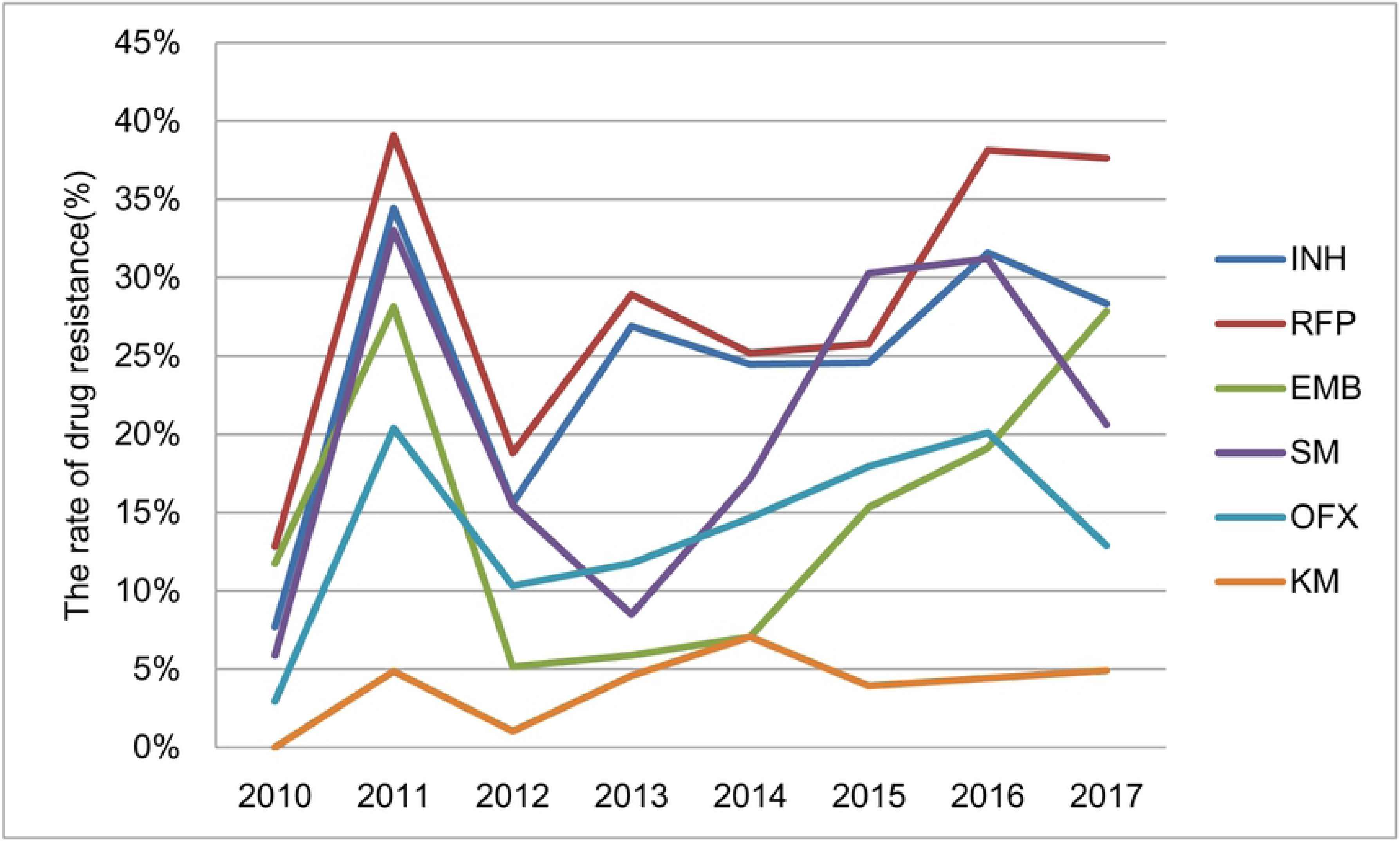
The drug resistance patterns in previously treated TB cases from 2010 to 2017.

In 2011, some drug resistance rates had a peak in previously treated TB cases, which were higher than the rates in 2010 and 2012(Fig 7). From 2010 to 2011, some drug resistance rates have increased significantly, including INH (χ^2^ trend=10.72, *P*=0.001), RFP (χ^2^ trend=9.51, *P*=0.002), SM (χ^2^ trend=9.64, *P*=0.002), and OFX (χ^2^ trend=5.73, *P*=0.02). From 2011 to 2012, some drug resistance rates have declined significantly, including INH (χ^2^ trend=16.16, *P*=5.81× 10^−5^), RFP (χ^2^ trend=16.95, *P*=3.84×10^−5^), EMB (χ^2^ trend=18.64, *P*=1.58×10^−5^), and SM (χ^2^ trend=8.27, *P*=0.004).

### Risk factors associated with MDR/RR-TB

The risk factors associated with MDR/RR-TB were analyzed (Table 1). Some groups of TB cases were more likely to have MDR/RR-TB in univariate analysis, including male (crude odds ratio [cOR], 1.4; 95% confidence Interval [CI], 1.26-1.55), farmer (cOR, 2.98; 95% CI, 2.7-3.28), TB cases aged over-60 years (cOR, 3.32; 95% CI, 2.2-5.02), previously treated TB cases (cOR, 7.63; 95% CI, 6.81-8.55), TB cases in Urban Districts (cOR, 0.33; 95% CI, 0.26-0.42), TB cases in New Urban Development Districts (cOR, 1.95; 95% CI, 1.51-2.53), TB cases in Northeast Districts (cOR, 2.7; 95% CI, 2.1-3.48).

**Table 1.**
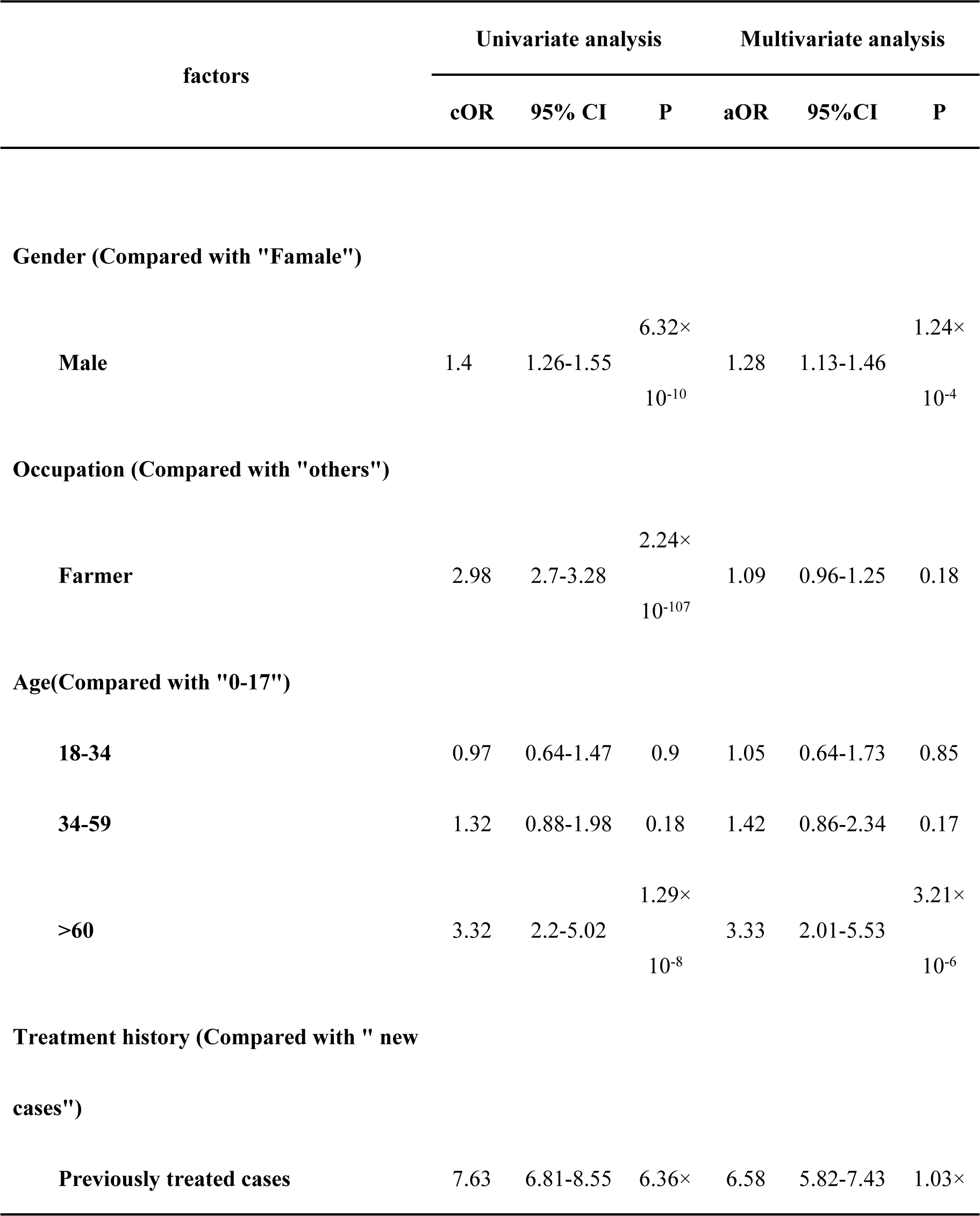
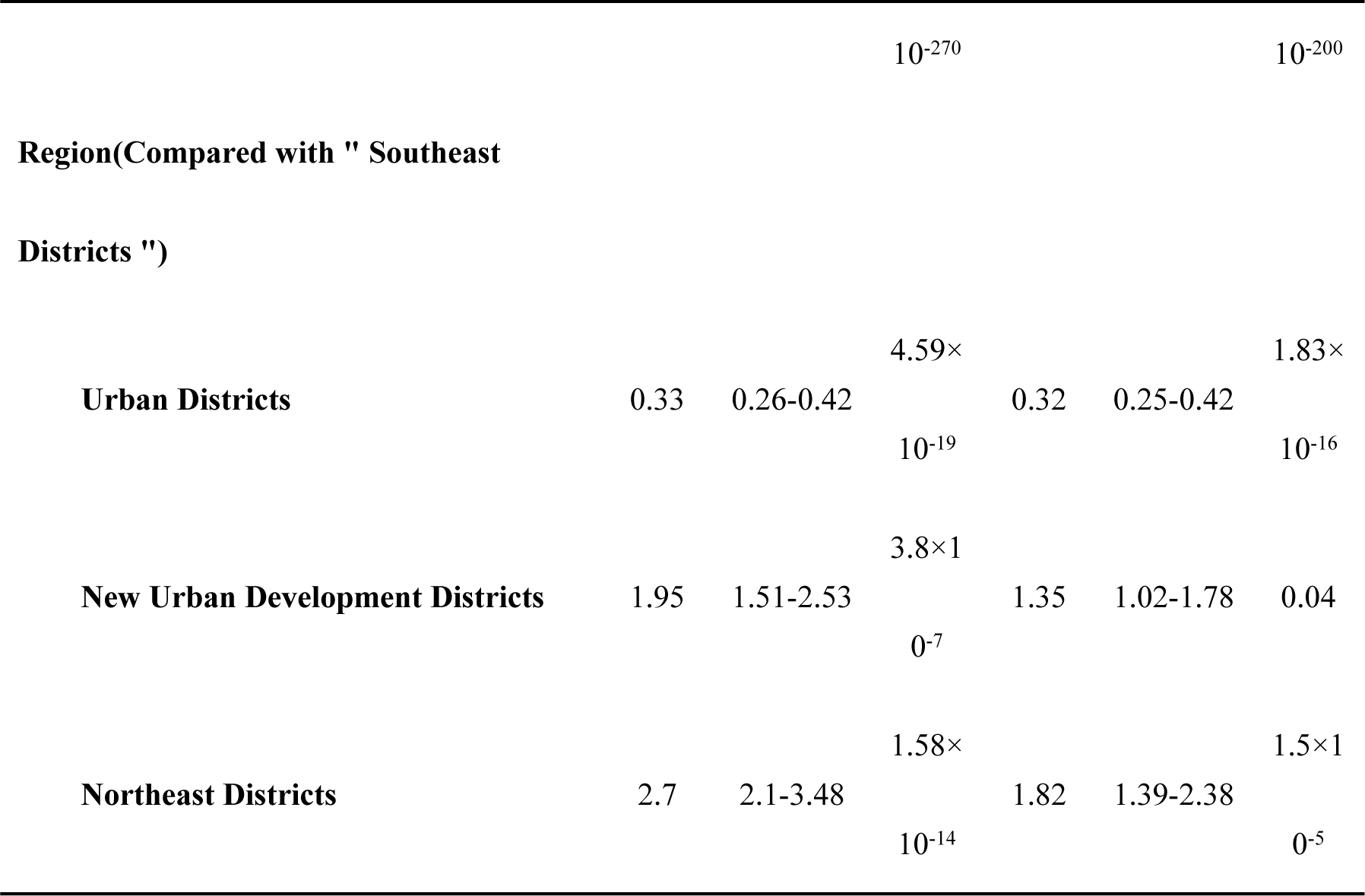
Risk factors associated with MDR/RR-TB.

| factors | Univariate analysis |  |  | Multivariate analysis |  |  |
| --- | --- | --- | --- | --- | --- | --- |
|  | cOR | 95% CI | P | aOR | 95%CI | P |
| **Gender (Compared with "Female")** |  |  |  |  |  |  |
| **Male** | 1.4 | 1.26-1.55 | 6.32×<br>10<sup>-10</sup> | 1.28 | 1.13-1.46 | 1.24×<br>10<sup>-4</sup> |
| **Occupation (Compared with "others")** |  |  |  |  |  |  |
| **Farmer** | 2.98 | 2.7-3.28 | 2.24×<br>10<sup>-107</sup> | 1.09 | 0.96-1.25 | 0.18 |
| **Age(Compared with "0-17")** |  |  |  |  |  |  |
| **18-34** | 0.97 | 0.64-1.47 | 0.9 | 1.05 | 0.64-1.73 | 0.85 |
| **34-59** | 1.32 | 0.88-1.98 | 0.18 | 1.42 | 0.86-2.34 | 0.17 |
| **>60** | 3.32 | 2.2-5.02 | 1.29×<br>10<sup>-8</sup> | 3.33 | 2.01-5.53 | 3.21×<br>10<sup>-6</sup> |
| **Treatment history (Compared with " new cases")** |  |  |  |  |  |  |
| **Previously treated cases** | 7.63 | 6.81-8.55 | 6.36× | 6.58 | 5.82-7.43 | 1.03× |
| | | | $10^{-270}$ | | | $10^{-200}$ |
| **Region(Compared with " Southeast** |  |  |  |  |  |  |
| **Districts ")** |  |  |  |  |  |  |
| **Urban Districts** | 0.33 | 0.26-0.42 | $4.59 \times 10^{-19}$ | 0.32 | 0.25-0.42 | $1.83 \times 10^{-16}$ |
| **New Urban Development Districts** | 1.95 | 1.51-2.53 | $3.8 \times 10^{-7}$ | 1.35 | 1.02-1.78 | 0.04 |
| **Northeast Districts** | 2.7 | 2.1-3.48 | $1.58 \times 10^{-14}$ | 1.82 | 1.39-2.38 | $1.5 \times 10^{-5}$ |

In multivariate analysis, MDR/RR-TB notification was not significantly associated with occupation (adjusted odds ratio [aOR], 1.09; 95% CI, 0.96-1.25). Other risk factors associated with MDR/RR-TB in univariate analysis were still correlated in multivariate analysis.

## Discussion

The MDR/RR-TB notification rate has increased significantly since 2010 in Chongqing. There were several reasons for this increase of MDR/RR-TB notification rate. One was that the foundation of MDR/RR-TB control system remained weak before 2009. The detection and treatment of MDR/RR-TB was not included in TB control plan of Chongqing in that period, and the MDR/RR-TB cases were not recorded in the TB surveillance system before 2010. This has contributed to the low MDR/RR-TB notification rate in early stage. Change began in 2009 by a project supported by the Bill and Melinda Gates Foundation [4]. This project aimed to improve MDR/RR-TB diagnosis and treatment in some counties and districts in 2011, including the following measures: the introduction of rapid molecular diagnosis, promotion of investment by local medical insurance and the foundation covering 90% of the medical cost, standard MDR/RR-TB treatment regimen and management, improved MDR/RR-TB detection, and establishment of cooperative mechanism between hospitals and CDC. Driven by this project, government funding for MDR/RR-TB has been rising year by year, and a sustainable financing mechanism has been established. After this project, the reimbursement rate of medical insurance for MDR/RR-TB has been increased to 90% with inpatient and outpatient care. The MDR/RR-TB screening has been financed by government since 2012, and rapid molecular diagnosis equipment was distributed to some laboratories. The scope of MDR/RR-TB screening has been expanded from five high-risk groups to all new TB cases and five high-risk groups. The laboratory diagnostic ability has been improved with continual training and supervision. The MDR/RR-TB reporting has been required in the electronic national TB surveillance system since 2010. With these improved measures, more and more MDR/RR-TB cases were notified. In 2017, the MDR/RR-TB notification rate was 2.09 cases per 100,000 population. But there still remained a big detection gap between the number of notified MDR/RR-TB cases and the estimate number of MDR/RR-TB cases among all notified TB cases. Screening should be further strengthened with more investment by government.

In Chongqing, the MDR/RR-TB notification rate of male was significantly higher than the rate of female, and male was more likely to develop MDR/RR-TB than female according to the correlation analysis. But gender impact on MDR/RR-TB was reported differently in different regions. There was no association with gender in several studies [8-13]. In some regions of China, studies showed that female was more likely to develop MDR/RR-TB [14, 15]. Socioeconomic condition could be one of the reasons for this difference. Chongqing was a city with a large poor rural population and many underdeveloped regions. Many men needed to go out to developed regions to work and support their families. Inadequate treatment and irregular management were more likely to happen in these cases with TB, which were important risk factors leading to MDR/RR-TB [16-18].

The MDR/RR-TB notification rate of over-60-year age group was not the highest in the four age groups. But our correlation analysis based on screening data showed that only over-60-year age group was the risk factor associated with MDR/RR-TB. There may be a contradiction between the two sets. This contradiction may indicate that the MDR/RR-TB notification in over-60-year age group was under-reported. Other study also showed that aging was a risk factor of MDR/RR-TB [11]. The correlation between aging and MDR/RR-TB might be due to socioeconomic status and common diseases of the elderly, like diabetes [19, 20]. The screening for MDR/RR-TB in elder TB cases should be strengthened.

The MDR/RR-TB notification rate of farmers was higher than the rate of all other occupations and increased to 4.11 cases per 100,000 population in 2017. Our study showed that the occupation of farmer was associated with MDR/RR-TB, and some studies in other regions of China also showed that farmer was a risk factor [21]. In Chongqing, farmer was a poor occupation. Although the income of farmer has gone up rapidly in recent years, the socioeconomic status of them was still low. According to Chongqing Statistical Yearbook, the annual income of farmers was less than half of urban residents. MDR/RR-TB were closely associated with poverty [22-24]. Medical services for MDR/RR-TB were often poorly accessible in rural areas [25].

The MDR/RR-TB notification rate of Urban Districts has been significantly higher than other three regions. But the region of Urban Districts was a protective factor in the correlation analysis for regional disparity. This was another contradiction in our study which may indicate that the detection of MDR/RR-TB was underestimated in other three regions. According to Chongqing Statistical Yearbook, the GDP per capita of Urban Districts was about two times higher than the level of other three regions from 2010 to 2017. The two largest prefectural designated hospitals, the provincial reference laboratory and provincial TB prevention and control institution were all located in Urban Districts. The lower socioeconomic status and weakness in health resources may lead to the under notification of MDR/RR-TB in other three regions.

Great progress has been made in the treatment of MDR/RR-TB. The treatment success rate increased significantly from 20% in 2011 cohort to 51.81% in 2015 cohort. The treatment success rate was 41% in 2014 cohort for whole China [2]. Cooperation between hospitals and CDC has proved successful. But 21.36% of notified cases were not involved in treatment, and 20.06% of them had not treatment outcome information reported. Even if medical insurance has covered 90% of the medical cost of MDR/RR-TB treatment, some poor cases still could not afford the out-of-pocket expenses, and were not involved in treatment because of poverty [26]. The information loss of treatment outcome indicated that cooperation between hospitals and CDC needed to be strengthened in the fields of information delivery and patient management.

From 2010 to 2017, the drug resistance rate for RFP in new TB cases was 6.23%, and this rate in previously treated TB cases was 32.7% in Chongqing. This rate was only 4.1% in new TB cases and 19% in previously treated TB cases globally [2]. Chongqing is facing high prevalence of MDR/RR-TB, especially in previously treated TB cases. The drug resistance rate for RFP increased significantly from 3.07% in 2010 to 15.53% in 2017. There were two possible reasons. One was that screening in high-risk groups has been effectively implemented in recent years, and more and more MDR/RR-TB cases were notified. A significant improvement in laboratory diagnostic capacity may be the other one. In 2011, some drug resistance rates had a peak in previously treated TB cases. In this year, some counties and districts screened MDR/RR-TB in sputum smear positive TB cases systematically supported by the Bill and Melinda Gates Foundation. Many MDR/RR-TB cases accumulated over the years have been notified, which led to this peak. In 2017, the drug resistance rate for RFP in new TB cases has declined significantly, and this was mainly due to the expansion of screening in 2017. All new TB cases including sputum smear negative have been required to be screened since 2017. The new screening policy has led to this drug resistance rate decrease in new TB cases.

This study has limitations. The epidemiological survey for MDR/RR-TB has not been implemented in recent years, so the routine surveillance data have been analyzed in our study. Only two second-line drug susceptibility results were recorded in the electronic surveillance system. The population could only be divided into four age groups according to Chongqing Statistical Yearbook.

In conclusion, MDR/RR-TB control over the years has been effective in Chongqing, and more and more MDR/RR-TB cases have been notified and treated. But the prevalence of MDR/RR-TB is still high, and facing the challenges including detection and treatment gaps, the regional disparity due to socioeconomic status, the under-notification in elderly people, and the high notification rate in farmers. Sustained government financing and policy support should be guaranteed to ensure universal access to effective MDR/RR-TB medical care.

